# Physical sensorimotor contingencies facilitate learning in monkeys

**DOI:** 10.64898/2026.11.01.763380

**Authors:** Su-Jin An, Jack Gabel, Mehrdad Jazayeri, Hansem Sohn

**Affiliations:** Department of Biomedical Engineering, Sungkyunkwan University (SKKU), Suwon, Republic of Korea; Center for Neuroscience Imaging Research, Sungkyunkwan University (SKKU), Suwon, Republic of Korea; McGovern Institute for Brain Research, Massachusetts Institute of Technology, Cambridge, Massachusetts 02139, USA; Department of Brain & Cognitive Sciences, Massachusetts Institute of Technology, Cambridge, Massachusetts 02139, USA; Howard Hughes Medical Institute, Massachusetts Institute of Technology, Cambridge, USA; Department of Brain Science and Engineering, Sungkyunkwan University (SKKU), Suwon, Republic of Korea; Department of Intelligent Precision Healthcare Convergence, Sungkyunkwan University (SKKU), Suwon, Republic of Korea

## Abstract

Animal studies in neuroscience routinely use virtual tasks that deviate significantly from the animals’ natural behavioral repertoire. This choice is often necessary for conducting careful and controlled experimentation. However, this approach raises a fundamental question: do virtual tasks expose neural computations and mechanisms associated with naturalistic tasks? In this study, we explore how naturalistic settings influence learning timescale and generalization behavior in rhesus monkeys. By training animals in a maze-solving task and comparing their behavior across two environments, we observe significant differences. Specifically, we found that a realistic setup using physical objects enabled learning that was orders of magnitude faster than the learning timescale observed in a comparable computerized virtual environment. Furthermore, we noted limited transfer of performance between the physical and virtual settings. These findings underscore the need for understanding how task environments shape learning and generalization, with potential implications for the design of future neuroscience and AI experiments.

## Main text

Learning efficiency and generalization capacity have been the focus of extensive research in psychology, neuroscience, and recently in AI. A key open question is when biological or artificial agents learn efficiently and generalize flexibly, and what factors influence these processes. In this context, we examine physical reasoning as a canonical cognitive domain, which has emerged as a core ingredient of ‘embodied approaches’ in AI. Previous studies in comparative psychology have shown that non-human primates possess inductive biases for constructing internal models of objects in the physical world—for instance, young chimpanzees use a simulation strategy to navigate a maze-solving task^1^. This capacity, often referred to as intuitive physics, is thought to benefit from inductive biases that are shared across various phylogenetic and ontogenetic stages^2^. Furthermore, neuroscience experiments have shown that neural activity in the frontal and temporal cortices exhibit features consistent with internal simulations of virtual objects^3,4^.

In this study, we compared monkeys’ learning and generalization across two task environments: a virtual, screen-based computerized maze and a physical, naturalistic maze with real objects (Figure 1). We first report the results from two monkeys (G and H) and then describe data from the other monkey (B) who underwent a different training curriculum and maze configurations. In the virtual maze condition (Figure 1A), animals were asked to predict the ball’s exit position and maneuver a joystick to catch it, akin to the game Pinball (Supplementary Figure 1). The training comprised four task conditions where we systematically manipulated maze configurations. In the first condition, the maze featured two straight paths. Both monkeys quickly learned this condition and surpassed chance performance (50% correct; Monkey H: after 27 trials, G: after 72 trials). Subsequently, we introduced a path with two turning points (“ㄣ”) in the second condition. One monkey (H) performed above chance level from the outset. The other monkey (G) initially chose the point that aligned vertically with the ball’s initial position. This apparent ‘gravity bias’^5^ disappeared after 3236 trials spread across six sessions. Next, in condition 3, we combined the two previous conditions by introducing two paths with two turnpoints. Initially, the monkeys’ performance was at chance level (H: 49.33%, G: 49.46%) and it took more than 20 daily sessions for them to attain 75% correct performance in this condition (H: 17,745 trials; G: 9,495 trials). This suggests that, in the virtual environment, animals did not flexibly generalize from their previous task learning. In the fourth final condition, the two paths overlapped, forming a junction where the ball continued straight along its originating path. Based on human intuitive physics, we expected that animals would learn the ball’s dynamics at the junction. Surprisingly, even after thousands of trials spanning nine sessions, both monkeys performed close to chance level (H: 58.76% with 1,940 trials; G: 35.5% with 7,385 trials). The animals showed a strong response bias toward one side (% left choice; H: 36.81%, G: 16.41%), suggesting that they resorted to a simple heuristic strategy. These results suggested that monkeys have a limited learning capacity for novel physical events in the virtual environment.

**Figure 1.**
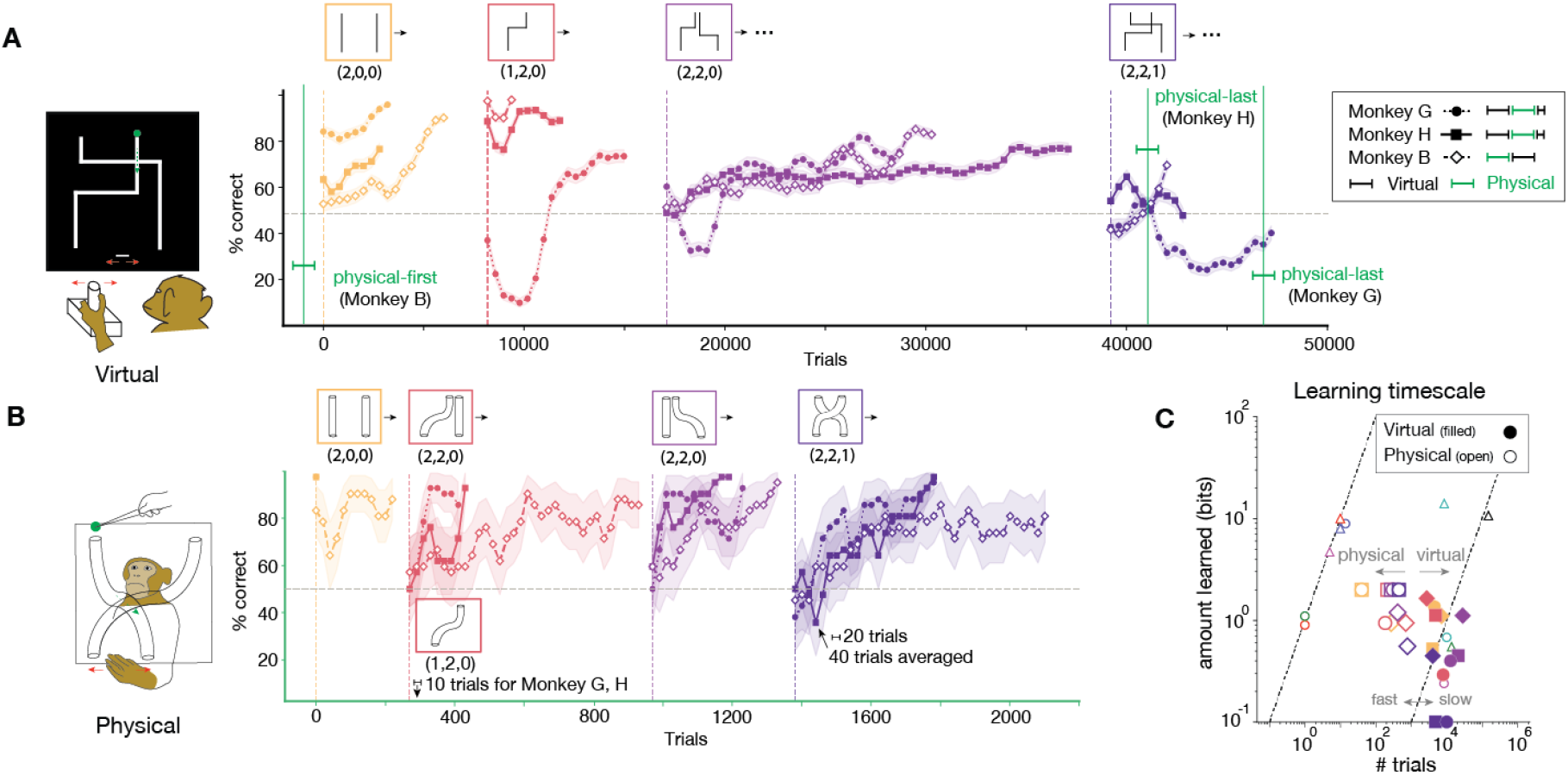
Monkeys learn to solve a maze faster in a naturalistic physical setting than in a virtual environment. (A) (left) Virtual maze-solving task: Monkeys control a paddle at the bottom of the screen by maneuvering a joystick to intercept a target ball, which navigates a maze from the top of the screen to the bottom. (right) Learning curves (moving average over 1000 trials, step size: 400 trials; shade: 95% binomial confidence interval with normal approximation [1.96 × SEM]). We systematically vary the maze configurations, changing the number of paths, turns, and junctions, with three numbers denoted in parentheses. The green vertical line and bracket indicate the period when we trained animals using a non-virtual (physical) version of the task. (B) (left) Naturalistic task setup. Monkeys interact with a maze composed of transparent tubes to reach and intercept a rewarding object using their hand. (right) Learning curves (moving average over 40 trials, step size: 20 trials; shade: 95% binomial confidence interval with normal approximation [1.96 × SEM]; green axes correspond to green brackets in panel (A)). For monkeys G and H, we use maze type (1 path, 2 turn, 0 junction) in the middle of training on maze type (2 paths, 2 turns, 0 junction) (arrow around the 100th trial) to reduce animals’ choice biases. (C) Learning rates. Animals learned faster and more effectively in the physical setting than in the virtual environment. We used an information-theoretic measure to quantify task complexity (“amount learned” on the ordinate) as a function of the number of reinforced trials^6^. Data from the present experiment are represented by filled markers (virtual) and open markers (physical), with marker shape indicating animal identity (G: circle, H: square, B: diamond; as in panel (A)). Other data points come from different tasks and different species^6^. For example, data near the left diagonal line (indicative of fast learning) are from rodent fear conditioning and navigation, whereas data near the right diagonal line (indicative of slow learning) are from macaque reaching, human golf, and rodent sensory two-alternative-forced-choice tasks.

Our findings in the virtual task were unexpected given previous neuroscience studies that demonstrated monkeys’ rapid generalization in other visuomotor domains ^3,4,7^. In the original virtual environment, monkey G and H failed to learn the maze junction condition, showed limited generalization across maze types, and exhibited a relatively slow timescale of learning. We next asked whether monkeys would exhibit faster learning in a more naturalistic setting involving real objects.

In the physical maze (Figure 1B), animals were trained to catch a grape by placing their hand below the appropriate exit of the maze. Initially, we trained the animals using a single straight tube and then introduced two straight paths, mimicking the virtual setup. The animals quickly achieved perfect performance, demonstrating rapid learning. Next, we replaced a straight tube with a curved one to create a hybrid maze consisting of one straight and one curved segment. Initially, performance was at chance level (H: 45%, G: 65%), but after brief training with the curved tube alone, performance rapidly surpassed chance levels within 20 trials in a session (H: 85%, G: 90%). This learning timescale was orders of magnitude faster than that observed in the virtual training (H: 17745 trials, G: 9495 trials; see Supplementary Figure 2 for detailed quantification). As a third condition, we tested whether the animals could generalize to a mirror-image version of the 1-straight-1-curved condition. Although immediate generalization (few-shot learning) was not observed, the animals learned to perform above chance within approximately 30 trials (H: 27 trials; G: 34 trials). Finally, we constructed a maze with a junction by positioning two tubes in a crossed fashion—a critical condition in which the animals had previously failed to learn in the virtual setting. Remarkably, they learned this junction condition in about 100 trials over four sessions (H: 110 trials, G: 95 trials). We further tested whether learning in this junction condition could transfer back to the virtual setup (Figure 1A). However, even after more than 1,500 trials (H: 2664 trials, G: 1840 trials), performance remained at chance level, suggesting a lack of transfer between the physical and virtual settings.

While these findings suggest a fundamental difference in monkeys’ intuitive physics under virtual and physical environments (Figure 1C), it is important to consider several confounding factors that can affect the animals’ learning timescales and generalization capacities (see Supplementary Information for details). Specifically, training order, trial repetition, and reward amount were different between the virtual and physical settings. First, monkeys were trained on the virtual task before transitioning to the physical task raising the possibility of meta-learning. Second, in the virtual task, a new maze configuration was presented after each correct trial, whereas the physical setting used a fixed maze for each condition. The virtual task therefore featured a wider array of maze configurations than the physical version. Third, the animals could be more motivated to learn in the physical setting due to the larger reward provided by a grape piece compared to the juice droplets used in the virtual setting. To control for these factors, we trained another monkey (B) in the counterbalanced order—first on the physical task and later on the virtual—with fixed maze configurations and matched reward setting.

Monkey B also learned the physical maze rapidly across all four conditions with comparable speed to monkey G and H (Figure 1B; training details in Supplementary Figure 3). For example, in the fourth condition with junction, B’s performance exceeded chance level after 83 trials (H: 110 trials, G: 95 trials). After transitioning to a virtual environment, nearly 30-fold more trials were required for monkey B to learn the four task conditions, showing much slower learning as in the other monkeys (virtual: total 27839 trials, physical: 975 trials; Figure 1C). This finding suggests that rapid learning in the physical maze cannot be attributed solely to meta-learning from prior virtual maze experience. For monkey B, we also revised the task design to better match the physical and virtual task: matching reward amount, repeating the same maze configuration after correct trial, and introducing a slant to the horizontal segment for gravity effect (see Supplementary Information). Nevertheless, we still observed slower learning rates in the virtual environment, suggesting that these factors were not critical in determining animals’ learning timescale. One notable difference was that unlike monkeys G and H, who never learned the junction condition in the original virtual environment, monkey B successfully learned to solve the junction maze in the revised virtual environment (Figure 1A; 2543 trials to reach performance above chance level). Because we simultaneously modified several factors for monkey B, we cannot isolate the contribution of individual factors and it remains to be further tested which task change underlies the different generalization capacity.

Our findings of slower learning in the virtual environment suggest that physical experience confers a substantial learning-rate advantage. An important next step is to identify the key factors that drive the difference in learning and generalization for physical reasoning. One prominent candidate is three-dimensional realism in the physical task, which we did not match in the 2D virtual stimuli. More careful control over these and related factors will shed light on the key dimensions that support intuitive physical reasoning across different contexts. Yet our study provided critical empirical evidence for the importance of task environments in engaging intuitive physics in primates, with implications for both designing laboratory experiments and interpreting behavioral and neural data, especially for questions related to learning.

## Supplementary figures

**Figure S1.**
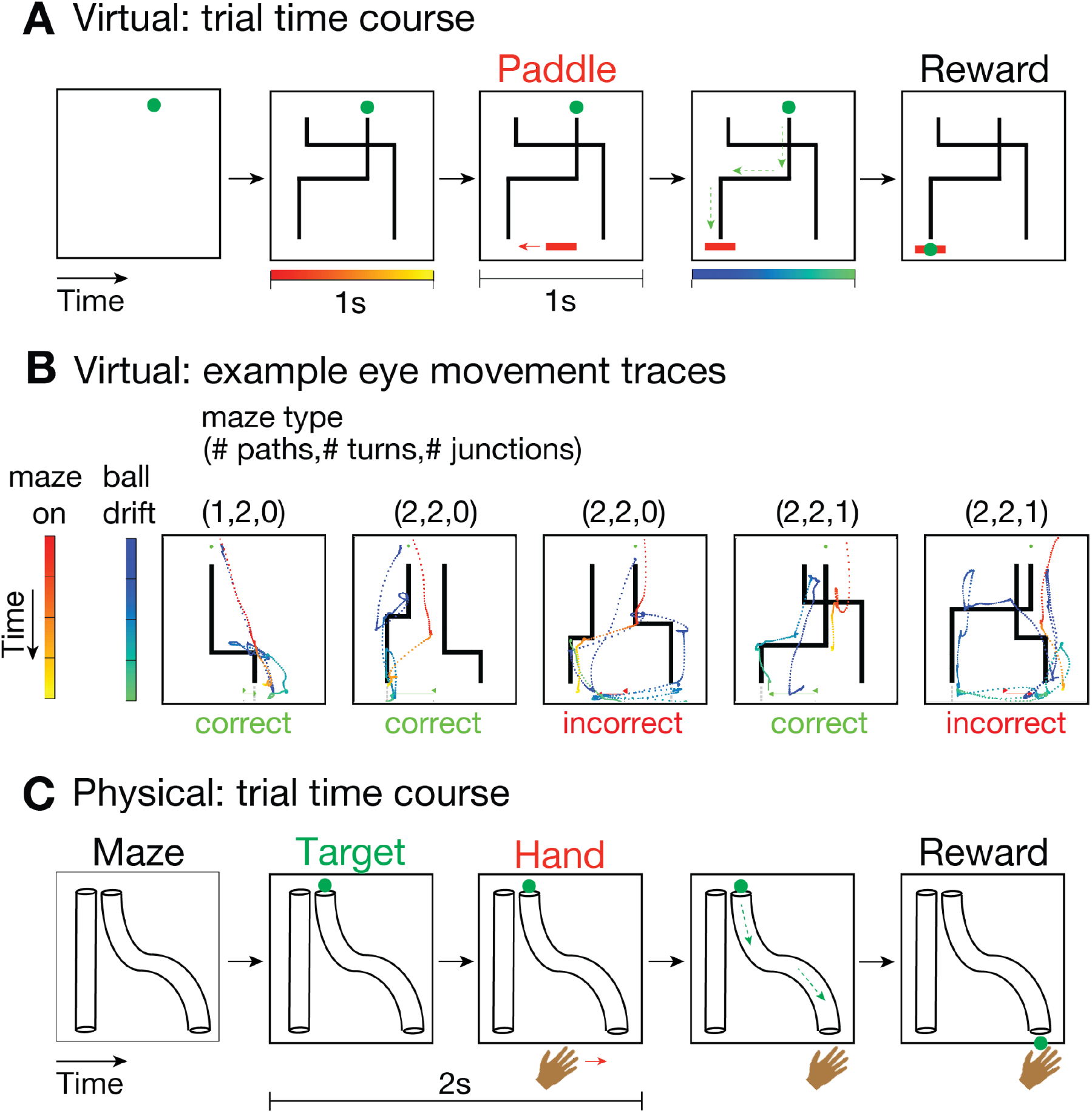
Trial time course of the maze-solving task in virtual and physical conditions, along with eye movement traces captured in the virtual setting. (A) Time course of an example trial in the virtual environment. Following the presentation of the target and maze (lasting for 1 second, denoted by a red-to-yellow line), the animals were given 1 second to maneuver the paddle before the target ball initiated its navigation through the maze (blue-to-green line). The animals received a reward if the paddle intercepted the ball. (B) Eye movement traces from five example trials in the virtual setup are shown. Eye positions are displayed during the maze-on epoch (red-to-yellow; 1 second) and the ball-drift epoch (blue-to-green; variable duration) superimposed on the maze stimulus. The arrows at the bottom center of each panel indicate the movement of the paddle at the onset and offset of the 1-second response epoch (green for correct, red for incorrect trials). These example trials vary in maze configurations, including the number of paths, turnpoints, and junctions. Notably, the animals fixated on the maze stimulus during the maze-on epoch before making a choice, suggesting that suboptimal task performance was not solely attributable to a lack of visual attention. (C) Time course of an example trial in the realistic physical environment. The target object is placed at the top entry, and the animals were given 2 seconds to reach for the target before it commenced traversing the maze. Monkeys were rewarded upon selecting the correct tube and successfully capturing the grape.

**Figure S2.**
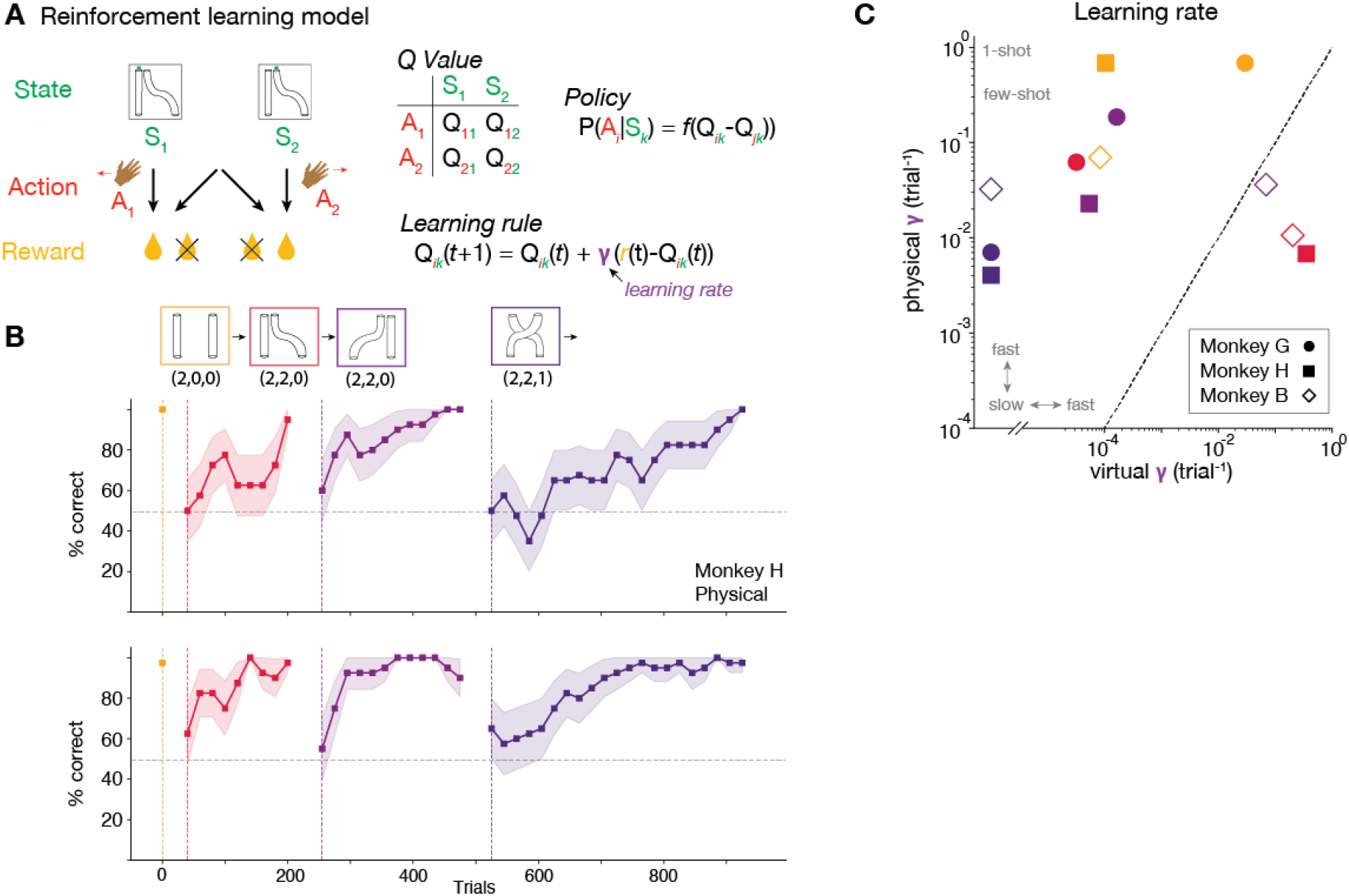
Learning rates, as measured by a reinforcement learning model ^8^, were faster in physical setup compared to the virtual environment. (A) A reinforcement learning model was used to measure the learning rate (i.e., the timescale of learning). (left) We defined the state space (initial target location), action space (animals’ choice), and corresponding reward based on the structure of the maze-solving task. (right) If we simplify the state and action spaces into two categories, there are four possible states for each state-action pair. The Q-learning model ^9^ was employed to update these values across trials using reward prediction error (*r*(*t*)-*Q*(*t*)) with a learning rate γ (as per the learning rule equation). The action is stochastically determined through a softmax function (*f*(.)) based on the value difference between two actions (as described in the policy equation). (B) The fitted Q-learning model successfully reproduced the animals’ behavior. (C) The learning rate was higher in the physical than the virtual setting. Different colors denote different task conditions (see panel (B)).

**Figure S3.**
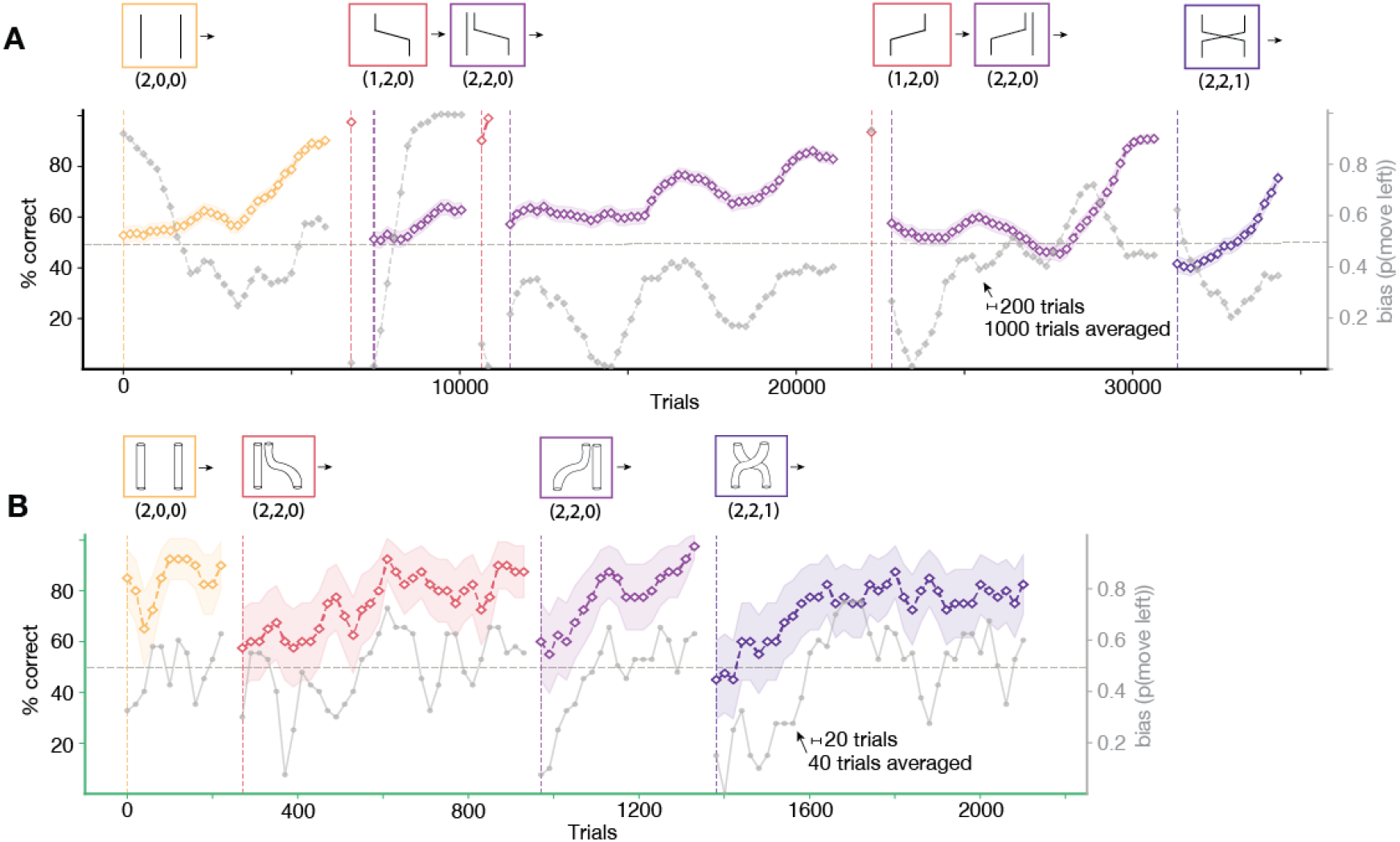
Full training trajectories, learning accuracy, and choice biases across virtual and physical maze conditions for Monkey B. (A) Virtual maze training. The complete training sequence across maze conditions is shown on the top. Learning curves were calculated using a moving average of 1000 trials with a step size of 200 trials. The left y-axis shows percentage correct, and the right y-axis shows choice bias, quantified as the probability of moving left, shown in gray. During the initial (2,2,0) training, a strong side bias emerged, prompting a return to the (1,2,0) condition before resuming (2,2,0) training. In each (2,2,0) condition, trials were repeated following incorrect or missed responses, and these repeated trials were excluded from accuracy calculations. In all conditions, trials in which the animal did not exit the response window (defined as a rectangle whose boundary extends about 48% of the distance from the starting position toward either maze exit) or did not respond were excluded from accuracy calculations. For the first (2,2,0) condition only, runs of five or more consecutive non-responses were treated as periods of disengagement and excluded, while all remaining trials—including isolated non-responses—were included in accuracy calculations. (B) Physical maze training. The complete training sequence across maze conditions is shown. Learning curves were calculated using a moving average of 40 trials with a step size of 20 trials. The left y-axis shows percentage correct, and the right y-axis shows choice bias, quantified as the probability of moving left, shown in gray. No repeated trials were used during physical maze training.

## Supplementary information

### Subjects

Three rhesus monkeys (Macaca mulatta) participated in the study: two males (G and H; 7 years old) and one female (B; 10 years old). All experimental procedures were approved by the Committee of Animal Care at the Massachusetts Institute of Technology (Cambridge, MA, USA) and Sungkyunkwan University (Suwon, Republic of Korea), and conformed to the guidelines of the National Institutes of Health.

### Virtual setup

Animals were comfortably seated in a dark, quiet room. Monkeys G and H were head-restrained to monitor their eye movements (Eyelink 1000, SR Research Ltd, Ontario, Canada), whereas monkey B was not head-restrained and eye movements were not recorded. All animals used a custom-made joystick to control a paddle on the screen. Prior to training on the maze-solving task, they underwent joystick training to achieve proficient control. Maze stimuli were presented on a 23-inch monitor (refresh rate: 60 Hz). We used MWorks software (https://mworks-project.org) to control visual stimuli and register animals’ hand movements, as well as eye movements for monkeys G and H. We procedurally generated the maze stimulus ^10^ to systematically vary the number of paths, turns, and junctions. For monkeys G and H, maze stimuli were randomly generated across trials, resulting in different maze shapes (i.e., different positions of paths, turns, and junctions). Following an incorrect response, the same maze configuration was repeated. For monkey B, the maze configuration was fixed within each condition, with no trial-by-trial variation, and matched the corresponding physical setting. Within the (2 paths,2 turns,0 junction) conditions, the ball’s entry position was repeated when the experimenter visually detected a strong response heuristic (e.g., a persistent side bias), with the frequency of repetition varying with the degree of the heuristic (Supplementary Figure 3). In the virtual environment used for monkeys G and H, turns consisted of instantaneous 90° changes in direction, whereas for monkey B, we revised the virtual maze so that each turn was connected by a diagonal segment (approximately 30° from horizontal) between the vertical segments before and after the turn, approximating the inclination of the physical maze’s tubes. Across all animals, the ball moved at a constant speed, with no friction or momentum modeled. Each animal completed approximately 600-1000 trials per daily session.

### Physical setup

Animals were seated comfortably in a brightly lit room, with chairs designed to allow them to reach toward the maze exit to obtain a reward (a grape for monkeys G and H; water droplet for monkey B). A transparent plexiglass plate attached to the chair provided a fronto-parallel view of the maze and also served as a platform for constructing the maze. The maze consisted of one or two transparent PVC tubes through which the animals could see a grape (monkeys G and H) or a wooden ball (monkey B) moving through the maze. Prior to the main maze-solving experiment, the animals underwent a familiarization stage using a straight tube to learn the task and reward contingencies. Within each of the four task conditions, the maze configurations remained fixed across trials. During each daily session, MATLAB or Python generated a random sequence of left- and right-side entry positions. On each trial, the experimenter (authors HS and SA) placed the object at the designated entry position. The animals were then given 2 s to predict which exit the object would emerge from by positioning their hand below the corresponding exit, after which the object was released into the maze. Their choices at the end of the 2-second response epoch were recorded manually. For monkey B, a water reward was delivered separately following a correct response. The entire experiment was video-recorded, and an independent observer (author JG) cross-validated the original manual coding for monkeys G and H (across-observer consistency; monkey H: 93.24%, monkey G: 92.93%), whereas monkey B’s responses were not independently cross-validated.

### Data Analysis

All analyses, including statistical tests, were performed using custom code in MATLAB (Mathworks, MA) and Python (version 3.13.2). To ensure a fair comparison between the virtual and physical settings and to equate the difficulty of responses between the joystick and hand, we discretized animals’ responses (i.e., paddle or hand position at the end of the response epoch) into a binary variable (left or right to the central midline). A response was considered correct if it matched the entry position, which was also binarized as left or right to the midline. This discretization allowed us to fit a reinforcement learning model (Supplementary figure 2) that is well-suited for handling discrete state/action spaces. To determine the trial number at which animals’ performance significantly exceeded chance levels (50% after discretizing responses), we employed a binomial test in a moving-window fashion. The window size was 20 trials for the first virtual condition (2 paths, 0 turnpoint, 0 junction; depicted by orange lines in Figure 1A) and for all physical conditions (Figure 1B). For the other condition in the virtual setting, a larger window size of 300 trials was used to account for its slow learning rate.

### Details of transfer test

After completing physical training, we tested whether monkeys G and H could transfer their successful learning of the junction condition to the virtual setting. Simplifying the junction condition to involve two stimulus states (the grape’s left or right position at the top) and two response options (left or right for hand at the bottom), the two-curved-path condition can be described as an exclusive OR operation in elementary logic ^11–13^. We reasoned that the animals could transfer their learning to the virtual setup by abstracting the state space for stimuli and responses and applying the same logical rule across different settings without relying on detailed task features. However, over four daily sessions comprising more than 1500 trials (H: 2664 trials, G: 1840 trials), the animals’ performance remained at chance, indicating that the intuitive physics learning was dependent on the specific stimuli and response modalities. This lack of transfer suggests that the learned behavior did not generalize across the different stimulus and response modalities used in the two settings. Thus, for monkeys G and H, learning in the naturalistic physical setting did not generalize to the virtual setting with a screen and joystick. We speculate that a critical computational process that may not be easily amenable in the virtual setting is the abstraction of state spaces for sensory input and motor output via embodiment^S1^.

### Control experiments and limitations

To examine potential alternative explanations for the learning differences between the two settings, we conducted several control experiments for monkeys G and H after their virtual training. First, we confirmed that the worse performance in the virtual environment was not due to poor joystick control or lack of visual attention. The animals demonstrated proficient joystick maneuvering skills, successfully receiving rewards in over 90% of trials in the easiest virtual condition (two straight paths). For monkeys G and H, we also monitored eye movements to assess attention to the maze stimuli (Supplementary Figure 1). We next examined whether the reward amount contributed to the difference in learning. We quantified the grape reward as 1.92 ± 0.08 g (mean ± SEM across six sessions) and adjusted the juice reward accordingly in the virtual task. Despite matched reward amounts, neither monkey G nor H learned the junction condition within 1500 trials. We also tested whether trial-by-trial variation in maze configuration contributed to the difficulty of the virtual task. When the maze configuration was held fixed across trials, neither monkey G nor H learned the junction condition within the tested training period (H: 1347 trials; G: 808 trials).

Monkey B provided an additional control that addressed several differences between the two settings simultaneously. Monkey B had no prior virtual experience and was trained in the reverse order (physical-first; virtual-first in G and H), received the same type of reward in both settings, and was trained on the virtual maze with a fixed maze configuration. In addition, the virtual environment was revised to more closely match the physical task. Under these conditions, monkey B learned the physical junction condition after 83 trials and subsequently learned the junction condition in the revised virtual environment after 2543 trials (Supplementary Figure 3). These results provide additional evidence that the difference in learning cannot be readily attributed to training order, reward amount, or trial-by-trial variation in maze configuration alone.

However, because the revised virtual environment incorporated multiple changes simultaneously, the contribution of individual factors cannot be isolated. Moreover, several differences between the physical and virtual settings remained. These included the three-dimensionality and physical realism of the environment, direct sensorimotor interaction with physical objects, the potential involvement of social factors through interaction with the experimenter, and differences in the physical dynamics of the environment. In particular, the virtual environment did not fully reproduce real-world physical dynamics, including realistic gravitational effects on the ball. Therefore, although our control experiments address several straightforward alternative explanations, they cannot fully determine whether the observed learning advantage in the physical setting arises from any single factor or from the combination of these differences. Future studies that systematically manipulate environmental realism, physical dynamics, sensorimotor interaction, and social factors will be necessary to determine which aspects of the physical environment are critical for rapid learning and generalization.

## Acknowledgments

M.J. is supported by the Howard Hughes Medical Institute (HHMI), NIH (R35NS137370), the Simons Foundation, and the McGovern Institute. S.J.A. and H.S. were supported by Basic Science Research Program through the National Research Foundation of Korea (NRF) funded by the Ministry of Education (RS-2025-25431602). S.J.A. is supported by funding from Institute for Basic Science (IBS) (R015-D1, R015-D2). The authors thank Nicholas Watters for assistance in implementing the virtual version of the maze-solving task.

## Data and code availability

The source data generated in this study and the custom scripts used for data analysis will be available in a public repository at publication.

## Declaration of Generative AI and AI-assisted technologies in the writing process

During the preparation of this work the authors used ChatGPT in order to improve the readability. After using this tool, the authors reviewed and edited the content as needed and took full responsibility for the content of the publication.

